# Molecular features of premenopausal breast cancers in Latin American women: pilot results from the PRECAMA study

**DOI:** 10.1101/396218

**Authors:** Magali Olivier, Liacine Bouaoun, Stephanie Villar, Alexis Robitaille, Vincent Cahais, Adriana Heguy, Graham Byrnes, Florence Le Calvez-Kelm, Gabriela Torres-Mejía, Isabel Alvarado-Cabrero, Fazlollah Shahram Imani-Razavi, Gloria Inés Sánchez, Roberto Jaramillo, Carolina Porras, Ana Cecilia Rodriguez, Maria Luisa Garmendia, José Luis Soto, Isabelle Romieu, Peggy Porter, Jamie Guenthoer, Sabina Rinaldi, on behalf of the PRECAMA team

## Abstract

**Background:** In Latin America (LA), there is a high incidence rate of breast cancer (BC) in premenopausal women, and the genomic features of these BC remain unknown. Here, we aim to characterize the molecular features of BC in young LA women within the framework of the PRECAMA study, a multicenter population-based case-control study on breast cancer in premenopausal women.

**Methods:** Pathological tumor tissues were collected from incident cases from four LA countries. Immunohistochemistry (IHC) was performed centrally for ER, PR, HER2, Ki67, EGFR, CK5/6 and p53 protein markers. Targeted deep sequencing was done on genomic DNA extracted from formalin-fixed paraffin-embedded (FFPE) tumour tissues and their paired blood samples to screen for somatic mutations in eight genes frequently mutated in BC. A subset of samples was analyzed by exome sequencing to identify somatic mutational signatures.

**Results:** The majority of cases were positive for ER or PR (168/233; 72%) and there were 21% triple negative (TN) cases, mainly of basal type. Most tumors were positive for Ki67 (189/233; 81%). In 126 sequenced cases, *TP53* and *PIK3CA* were the most frequently mutated genes (32.5% and 21.4%, respectively), followed by *AKT1* (9.5%). *TP53* mutations were more frequent in HER2-enriched and TN IHC subtypes, while *PIK3CA/AKT1* mutations were more frequent in ER positive tumors, as expected. Interestingly, a higher proportion of G:C>T:A mutations was observed in *TP53* gene in PRECAMA cases compared to TCGA and METABRIC breast cancer series (27% vs 14%). Exome-wide mutational patterns in 10 TN cases revealed alterations in signaling transduction pathways and major contributions of mutational signatures caused by altered DNA repair pathways.

**Conclusions:** This pilot results on PRECAMA tumors gives a preview of the molecular features of premenopausal BC in LA. Although, the overall mutation burden was as expected from data in other populations, mutational patterns observed in *TP53* and exome-wide suggested possible differences in mutagenic processes giving rise to these tumors compared to other populations. Further omics analyses of a larger number of cases in the near future will allow investigating relationships between these molecular features and risk factors.

## Introduction

Breast cancer (BC) incidence is increasing sharply in countries in economic transition with a large number of cases in premenopausal women. In Latin America (LA), the frequency of BC in women younger than 45 years is close to twice the frequency seen in developed countries, an increase only partly explained by population age-structure [1]. Behavioural, reproductive and lifestyle factors typical of the Western populations are becoming more prevalent in LA and may play a role in the increased BC incidence in this population, but the reason for the sharp increase in premenopausal women remains to be established [2].

BC is a heterogeneous disease in terms of biology and outcome. It is clinically classified into four subtypes (Luminal A, Luminal B, HER2 positive, Triple-negative) based on the expression of the oestrogen receptor (ER), progesterone receptor (PR), HER2 receptor and the proliferation marker Ki67 [3]. More sophisticated classifications based on genomic and transcriptional analyses provide a better description of the tumor biology and outcome [4, 5]. The two most frequently somatically mutated genes in BC are *TP53* and *PIK3CA*. Mutations in *PIK3CA*, which renders cells dependent on *PI3K* pathway signalling, are the most common genetic abnormality identified in hormone receptor positive breast cancer, while mutations in the tumour suppressor gene *TP53* are more prevalent in *HER2*-enriched and triple-negative (TN) subtypes.

Genomic analyses can also provide information related to tumor aetiology. Indeed, somatic mutational signatures can reveal the contribution of specific mutational processes to the development of cancer. For example, *TP53* mutation patterns specific to exposure to exogenous mutagens have been reported in several cancer types [6], and at the genome-wide level, over 30 mutational signatures have been described in cancer tissues and some have been linked to endogenous mechanisms of mutagenesis or to exposure to human carcinogens [7, 8].

While BC genomic subtypes have been associated with different patient outcomes, how specific genomic alterations relate to risk factors or aetiology remains largely unknown. Moreover, knowledge of the genomic features of premenopausal BC, particularly in countries in economic transition, is limited. The PRECAMA study was initiated to investigate the molecular, pathological, and risk factor patterns of premenopausal BC in LA (http://precama.iarc.fr/). It is the largest case-control study conducted in four Latin American countries that systematically collects both, extensive information on life-style and risk factors, and different biological samples (tumor tissues, blood fractions and urines) according to standardized procedures. PRECAMA is thus a powerful framework for investigating relationships between BC tumor biology and aetiology.

Here, we investigate the tumor genomic features of premenopausal BC (preBC) in Latin American women using the first set of samples collected within the framework of the PRECAMA study.

## Materials and Methods

### Study population

Subjects were women age 20-45 years diagnosed with BC as part of the PRECAMA case-control study (http://precama.iarc.fr). Recruitment was conducted at major general or cancer dedicated hospitals in Chile, Colombia, Costa Rica and Mexico which cover populations with a wide range of socio-economic status. Women having a positive biopsy for BC were recruited prior to any treatment. Women were invited to a home or hospital visit during which a trained nurse presented the informed consent, collected biological samples and anthropometric measurements (height, weight, hip and waist circumferences) and administered a standardized questionnaire on clinical, reproductive, and life-style risk-factors. All participants gave written informed consent before enrolment, and the study protocols were approved by the institutional review boards of Chile (Oncologic Institute Foundation Arturo Lopez Pérez and National Cancer Institute), Colombia (Cancer Institute Las Americas and University of Antioquia), Costa Rica (Costa Rican Institute of Clinical Research (ICIC) and Center for Strategic Development and Information in Health and Social Security (CENDEISSS) of the Costa Rican Social Security Fund (CCSS)), Mexico (National Institute of Public Health and the Social Security Mexican Institute), and the International Agency for Research on Cancer (IARC).

### Biological specimens

Each study site applied common standardized protocols for specimen collection. Protocols have previously been developed and extensively used by IARC [9, 10], and subsequently fine-tuned based on a detailed review of the conditions at each center. Blood samples were obtained by venipuncture using vacutainers at recruitment and buffy coats were prepared and stored at −80°C within less than 6 hours after blood drawn. Buffy coats were shipped to IARC for genomic DNA extraction. Tumor samples were fixed in formalin and paraffin-embedded (FFPE) according to SOPs. Paraffin blocks and H&E sections were stored at the local pathology service facilities. Sections from tumor tissues were sent to Fred Hutchinson Cancer Research Center (FHCRC) for centralized immunohistochemistry (IHC) analyses and tumor DNA extraction.

### Pathology Review and IHC analyses

Histology sections from tumor biopsy obtained prior to any treatment were reviewed for histological diagnosis and grade, lympho-vascular invasion and stromal and lymphocyte response. IHC was conducted for ER (SP1, LabVision, Fremont CA), PR (PgR 636, DAKO, Denmark), HER2 (AO485, DAKO, Denmark), EGFR (31G7 Invitrogen, Camarillo CA), CK5/6 (D5/16 B4, DAKO, Denmark), p53 (Pab 1801, Calbiochem, La Jolla CA), Ki-67 (MIB-1, DAKO, Denmark) according to standardized and optimized protocols that included antigen retrieval when required. BCs were classified into subtypes according to ER, PR and HER2 IHC results. Triple negative (ER-, PR-, HER2-) were additionally subtyped using EGFR and CK5/6 staining to define basal-like cancers. ER and PR positivity were defined as staining score >1%, and Ki67 positivity as staining >14% as recommended by the St Gallen International Breast Cancer Conference [3].

### DNA extraction and sequencing

Tumor genomic DNA was extracted from 3 to 9 sections of 6um using the QIAamp DNA FFPE Tissue Kit (Qiagen) following manufacturer’s recommended protocol, with the following modification. The tissue was incubated in ATL buffer and proteinase K overnight at 56°C with agitation, with an additional 20 uL of proteinase K addition after the first four hours. Constitutive genomic DNA was isolated from buffy coats at IARC with the Autopure LS system (Qiagen) using the frozen buffy coat protocol and following manufacturer instructions. DNA was quantified by PicoGreen® (ThermoFisher Scientific).

For **targeted sequencing**, exonic regions of the selected gene panel (AKT1, CDH1, ERBB2, NOTCH1, PIK3CA, PTEN, RB1, TP53) were amplified from 80 ng of genomic DNA using GeneRead DNAseq Mix-n-Match Panel V2 (Qiagen) following manufacturer’s instructions. Libraries were prepared with NEBNext reagents (New England Biolab) following manufacturer’s instructions. Libraries were quantified by PicoGreen® (ThermoFisher Scientific), pooled in equal quantities and the library pool was quantified by Qubit fluorometer (ThermoFisher Scientific) and quality checked with the Bioanalyzer (Agilent Technologies). 800pM of the library pool was used for sequencing on a Ion Proton sequencer (Life Technologies) according to manufacturer’s instructions, aiming at a minimum of 100X coverage for blood DNA and 1000X coverage for tumor DNA. Tumor samples were processed in duplicates to control for artefactual mutations from FFPE fixation (see bioinformatics analyses below).

For **whole-exome sequencing** (WES), exonic regions and splice junctions of tumor-blood DNA samples pairs were captured using the SeqCap EZ MedExome kit (Roche Diagnostics France) following manufacturer’s instructions. This assay captures exonic regions covering 47Mb of protein-coding bases. Libraries were prepared with the KAPA Hyper Prep Kit (Roche Diagnostics France) following manufacturer’s instructions and sequenced by 150-base paired-end massively parallel sequencing on an Illumina HiSeq 4000 sequencer at the New York University Langone Medical Center according to manufacturer instructions.

### Bioinformatic analyses

Data from the Ion Proton were processed with the Ion Torrent built-in pipeline (TorrentSuite V4) to generate BAM files and variant calling was done with the built-in ITVC in the somatic mode and with a minimum allelic frequency threshold of 4%. Variants were annotated with Annovar and filtered to eliminate known SNPs (variants present in ExAC or 1000G databases at frequency >0.001) and sequencing artefacts using our MutSpec Galaxy package developed in-house [11]. Further manual checks of BAM files using IGV were done when appropriate. All non-synonymous mutations found in the targeted regions and present in both duplicates of tumor samples but not in any blood sample were retained for analysis.

Exome data from the HiSeq4000 were analysed with a pipeline developed in-house and based on standards tools for quality control and processing (FastQC 0.11.3, AdapterRemoval 2.1.7, BWA-MEM 0.7.15, Qualimap 2, GATK 3.5, Picard 1.131). Somatic variant calling was done on tumor-blood sample pairs with Strelka [12] using default parameters. Variant annotation and filtering was done as described above with MutSpec [11] and only somatic indels and SNVs in coding regions were retained and analysed. Pathway analysis of mutated genes was done with ConsensusPathDB (r32) using KEGG, Biocarta, Reactome and Wikipathways databases and a minimum of 3 overlapping genes and q-value <0.05 as settings [13]. To define cancer genes, we used COSMIC Cancer Gene Census (v82)[14], and genes identified as drivers in breast cancer in the IntOGen database (r2014.12)[15].

### Public data on somatic mutations in breast cancer

Data from TCGA breast and METABRIC genomic studies [16, 17] and from the IARC TP53 Database [18] were used as comparison datasets. Gene-specific mutation files (*AKT1*, *CDH1, ERBB2, NOTCH1, PIK3CA, PTEN, RB1, TP53* genes) and related clinical files for TCGA and METABRIC studies were retrieved from cBioportal [19, 20] in February 2017. MAF files from exome sequencing data of TCGA breast cancer cases were retrieved on 26 March 2015 *via* a https protocol at https://tcga-data.nci.nih.gov/tcgafiles/ftp_auth/distro_ftpusers/anonymous/tumor/. Gene-specific data from TCGA and Metabric were combined, including only cases with documented age and ER, PR and HER2 status, and stratified by age (under 45 or above 55 years old). For TCGA exome data, only data with documented age and ER, PR and HER2 status were selected resulting in a dataset of 453 samples including 96 preBC and 357 postBC. Version R18 of the somatic dataset of the IARC TP53 Database was used selecting for mutations reported in primary breast cancers in women <=45 years old and in studies using Sanger sequencing.

### Statistical analyses

For mutational signatures analyses, we used PRECAMA exome data (N= 12 samples) and TCGA exome data (N = 453). Mutations were classified into 96 types corresponding to the six possible base substitutions (C:G>A:T, C:G>G:C, C:G>T:A, T:A>A:T, T:A>C:G, T:A>G:C) and the 16 possible pairs of flanking nucleotides immediately 5’ and 3’. Mutational signatures in these samples were then extracted using the non-negative matrix factorization (NMF) algorithm implemented in the NMF R package [8, 21]. NMF decomposition identifies signatures and estimates their contributions to each sample. Six signatures were identified using the cophenetic correlation coefficient [20] as a measure of stability of the signatures. We calculated, the cosine similarity between the six extracted signatures and those published in the Catalogue of Somatic Mutations in Cancer (COSMIC) and in other original reports [21, 26, 27], as described elsewhere [21].

We wished to identify possible systematic differences of signature contributions both between IHC subtypes, by menopausal status and by study source (TCGA vs PRECAMA). Due to the small number of PRECAMA samples, we used 2000 permutations of samples to obtain an empirical distribution of the Kruskal-Wallis rank-sum statistic. This permutation test was applied to test for possible association of each signature with a) menopausal status; b) IHC subtype; c) Menopausal status stratified by IHC subtype; d) menopausal status adjusted for subtype by linear model; and e) study source, with partial adjustment for subtype (TN vs others) and also restricted to pre-menopausal samples.

All statistical analyses were performed using the R statistical software version 3.3.2. The statistical significance level was set to 0.05 without adjustment for multiple comparisons.

## Results

### IHC subtypes in PRECAMA tumors

In the first consecutive cases recruited in PRECAMA and for which tumor pathological evaluation has been completed (N=229), most BC cases were ER positive (72%) and 16% were positive for HER2 (**Table 1**). Using ER/PR/HER2 IHC subtyping, the majority of cases were Luminal A (58%), followed by TN (21), Luminal B (11%) and HER2 enriched-enriched (5%) (**Table 1**). The HER2+ rate overall was 16%. TN tumors were predominately basal-like (98%) (**S1 Table**). Proliferation status was assessed by Ki67 IHC. More than 80% (189/233) of cases had high Ki67 staining (staining >= 14%) with a median percentage of 31.6 (not shown). Ki67 positive cases were less frequent among Luminal A cases than other subtypes (71.6% compared to 88-95%).

**Table 1.** Sample classification by IHC results.

|  | Sample number<br>N (%) | Ki67 positive<br>N (%) |
| --- | --- | --- |
| **ER** |  |  |
| **Negative** | 65 (28%) | 63 (97%) |
| **Positive** | 168 (72%) | 126 (75%) |
| **PR** |  |  |
| **Negative** | 71 (30%) | 67 (94%) |
| **Positive** | 162 (70%) | 122 (75%) |
| **HER2** |  |  |
| **Negative** | 183 (79%) | 144 (79%) |
| **equivocal** | 13 (6%) | 12 (92%) |
| **Positive** | 37 (16%) | 33 (90%) |
| **Total** | 233 | 189 (81%) |
| **IHC SUBTYPE\*\*** |  |  |
| **Luminal A** | 134 (58%) | 96 (72%) |
| **Luminal B** | 26 (11%) | 23 (88%) |
| **HER2 Enriched** | 11 (5%) | 10 (91%) |
| **Triple Negative** | 48 (21%) | 47 (98%) |
| *Of basal type* | 45 (94%) | 42 (93%) |
| **Undetermined\*\*\*** | 14 (6%) | 13 (93%) |

\* 20 cases were excluded due to no invasive tumor present or insufficient tissue for testing.
\*\* Tumor Subtype definitions: Luminal A: ER+/HER2-; Luminal B: ER+/HER2+; HER2 Enriched: ER-/HER2+; Triple Negative: ER-/PR-/HER2-; TN of basal type: EGFR and/or CK5/6 positive.
\*\*\* 14 cases were not assigned a subtype: 13 cases have equivocal HER2 results and no confirmatory FISH; 1 case is ER-/PR+/HER2- with a weak PR positivity.

### Somatic mutations in premenopausal BC

Tumor genomic DNA was extracted from FFPE tissue sections prepared in each collecting Centre according to a standardized protocol. More than 250 ng of DNA were obtained for 75% of the samples with a median yield of 994 ng. Limiting amount of DNA (<100 ng) was obtained for 12% (21/172) of the samples. Targeted deep sequencing of a panel of 8 genes frequently mutated in BC (*AKT1, CDH1, ERBB2, NOTCH1, PIK3CA, PTEN, RB1, TP53*) was successfully performed on 126 cases for which over 250 ng of tumor genomic DNA was available. Tumor DNA and matched blood DNA were sequenced at minimum coverages of 1000X and 100X, respectively. To control for potential artefacts due to formalin fixation, FFPE tumor samples were sequenced in duplicates and only mutations detected in both duplicates were considered (see Materials and Methods). Potentially deleterious somatic mutations (affecting splicing, indels, nonsense, stoploss, and non-synonymous substitutions) in the 8-gene panel were found in 63.5% (80/126) of samples. *TP53* was the most frequently mutated gene (32.5%), followed by *PIK3CA* (21.4%) *and AKT1* (9.5%), while other genes were mutated in less than 5% of samples. Overall, this distribution was similar to the one obtained in preBC women from TCGA/METABRIC datasets (top mutated genes are *TP53* and *PIK3CA*), although the prevalence of *TP53, PIK3CA, AKT1 and RB1* genes mutation prevalence were significantly different (**Fig 1A**). There were fewer cases with *TP53* and *PIK3CA* mutations and more cases with *AKT1* and *RB1* mutations (p<0.05). These differences may be explained in part by the lower proportion of TN and HER2-enriched cases (known to carry *TP53* and both *TP53* and *PIK3CA* mutations respectively [22]) and higher proportion of Luminal A cases (known to carry *AKT1* or *PIK3CA* mutations [22]) in PRECAMA samples compared to the TCGA/METABRIC datasets (see **S1 Fig**). Nonetheless, the relative proportion of *PIK3CA* versus *AKT1* mutations in Luminal A cases was different between PRECAMA and TCGA/METABRIC datasets, with a higher proportion of *AKT1* mutated cases in PRECAMA versus TCGA/METABRIC (14% *AKT1* and 23% *PIK3CA* mutations *vs* 4% *AKT1* and 54% *PIK3CA* mutations, respectively, although this was not statistically significant). Thus, while the *PIK3CA*/*AKT1* pathway is mutated at expected rates in the Luminal A PRECAMA tumors, *AKT1* mutations may be favoured over *PIK3CA* mutations compared to other published sets, a result that will need to be confirmed in a larger sample set. *PIK3CA* and *AKT1* mutations were at classical hotspots (p.H1047R, p.E542K, p.E545K for *PIK3CA* and p.E17K for *AKT1*), and *TP53* mutations were mostly missense substitutions that spread across the coding sequence (**S1 Table**). The relationship between IHC subtypes and mutated genes was as expected from previous studies (**Fig 1B**). *TP53, RB1* and *PTEN* mutated samples were more prevalent in TN samples, while *AKT1* or *CDH1* mutated samples were of Luminal A subtype. The majority of samples with no mutation in the tested genes were Luminal A cases (34/46; 74%).

**Fig 1.**
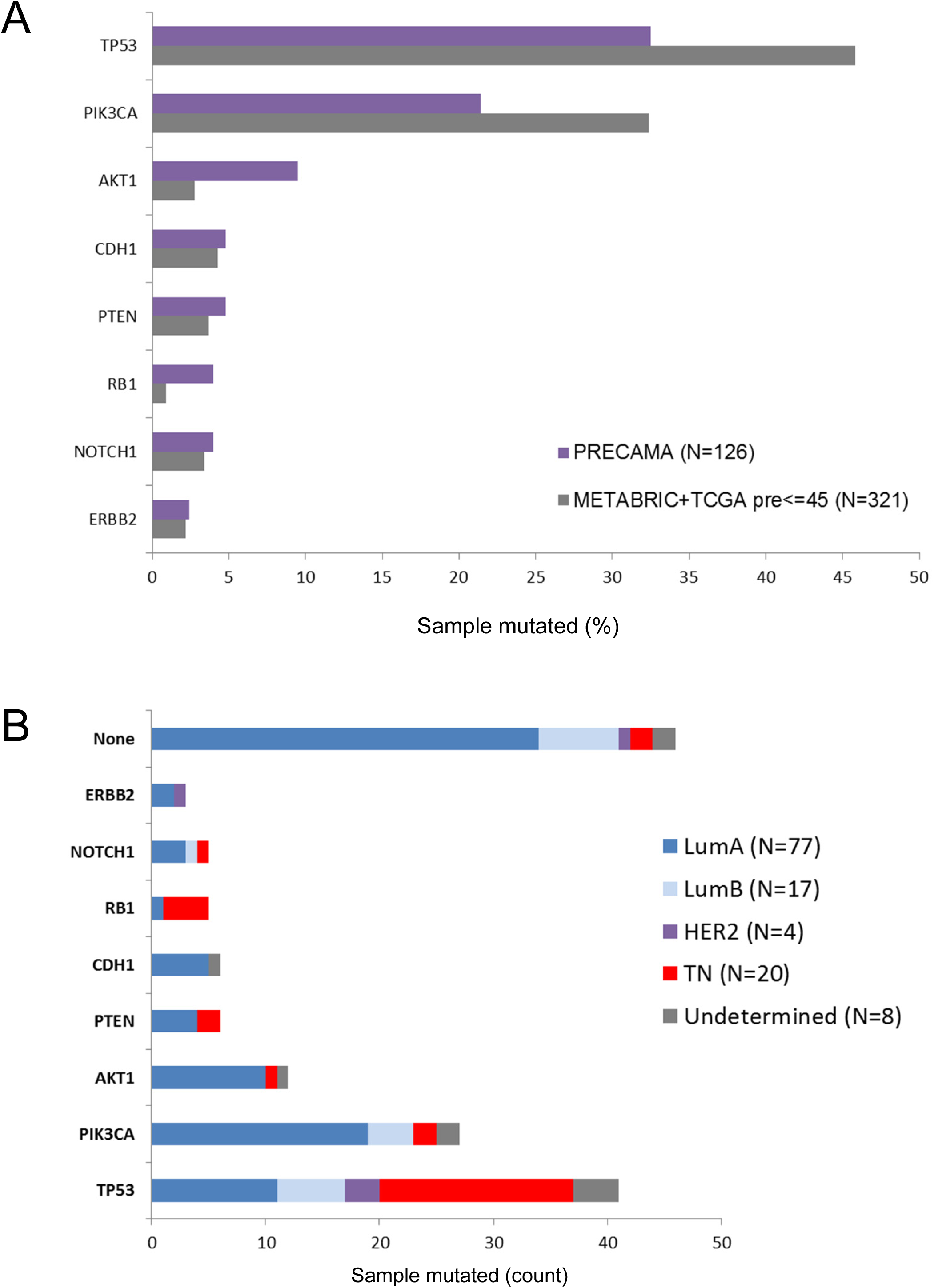
Mutation occurrences in eight BC genes. (A) Gene mutation frequencies in PRECAMA samples are compared to those observed in a dataset of premenopausal women selected from the TCGA and METABRIC series [16, 17]. (B) IHC subtype distributions of samples according to their mutation status. Luminal A: ER+/HER2-; Luminal B: ER+/HER2+; HER2 Enriched: ER-/HER2+; Triple Negative: ER-/PR-/HER2-.

Twenty one tumors had mutations in more than one gene. One case was of HER2-enriched subtype and had mutations in *TP53* and *ERBB2*. Three cases were of Luminal B subtype and had mutations in *PIK3CA* and *TP53* or *CDH1*. Seven cases were TN and had mutations in *TP53* combined with *RB1* (3 cases) or *PTEN* (2 cases) or *PIK3CA* (1 case) or *NOTCH1* (1 case). Ten cases were of Luminal A subtype and had mutations in *TP53* and *PIK3CA* (4 cases), in *TP53* and *AKT1* (3 cases), or other gene combinations. Mutation details are provided in **S1 Table**.

In a subset of 12 samples (two Luminal A and ten TN cases selected randomly) analyzed with the 8-genes panel, we also performed whole exome sequencing. With a median coverage of 200X in tumor DNA and 80X in blood DNA and over 99.5% of mapped reads (see **S2 Table**), we identified 2634 somatic mutations in coding regions, including 2128 non-synonymous SNVs and indels (see **S3 Table**). All mutations found by targeted sequencing in the 8-gene panel were confirmed in the exome analysis. There was an average of 3.9 non-synonymous SNVs and indels per MB, with 2 samples carrying more than 6 mutations per MB (**Fig 2A, top panel**). The top mutated genes included four cancer genes (*TP53*, *RB1, PIK3CA* and *AHNAK*) and several large sized genes such as mucin genes and *TTN* gene (**Fig 2A, middle panels**). *AHNAK* has recently been described as a novel tumor suppressor gene in breast cancer, especially in TN subtype, acting *via* different signaling pathways such as AKT/MAPK or TGFβ [23, 24]. *AHNAK*, like *RB1* mutations were all in TN cases. However, the impact of *AHNAK* mutations on protein function is unknown as *AHNAK* is a large size gene and 3/4 of the mutations were predicted as benign by Polyphen-2 [25]. In TN cases (N=10), there were 92 cancer genes mutated, dominated by *TP53* which was mutated in all samples, and with 14 other cancer genes mutated in more than one sample (**Fig 2B**). Pathway enrichment analysis of potential driver mutations in these TN samples (**S4 Table**) showed enrichment for several growth factor signaling pathways and for pathways involved in insulin receptor signaling, telomere maintenance, transmembrane transport of small molecules, G1 checkpoint or O-glycan biosynthesis (**Fig 2C and S5 Table**).

**Fig 2.**
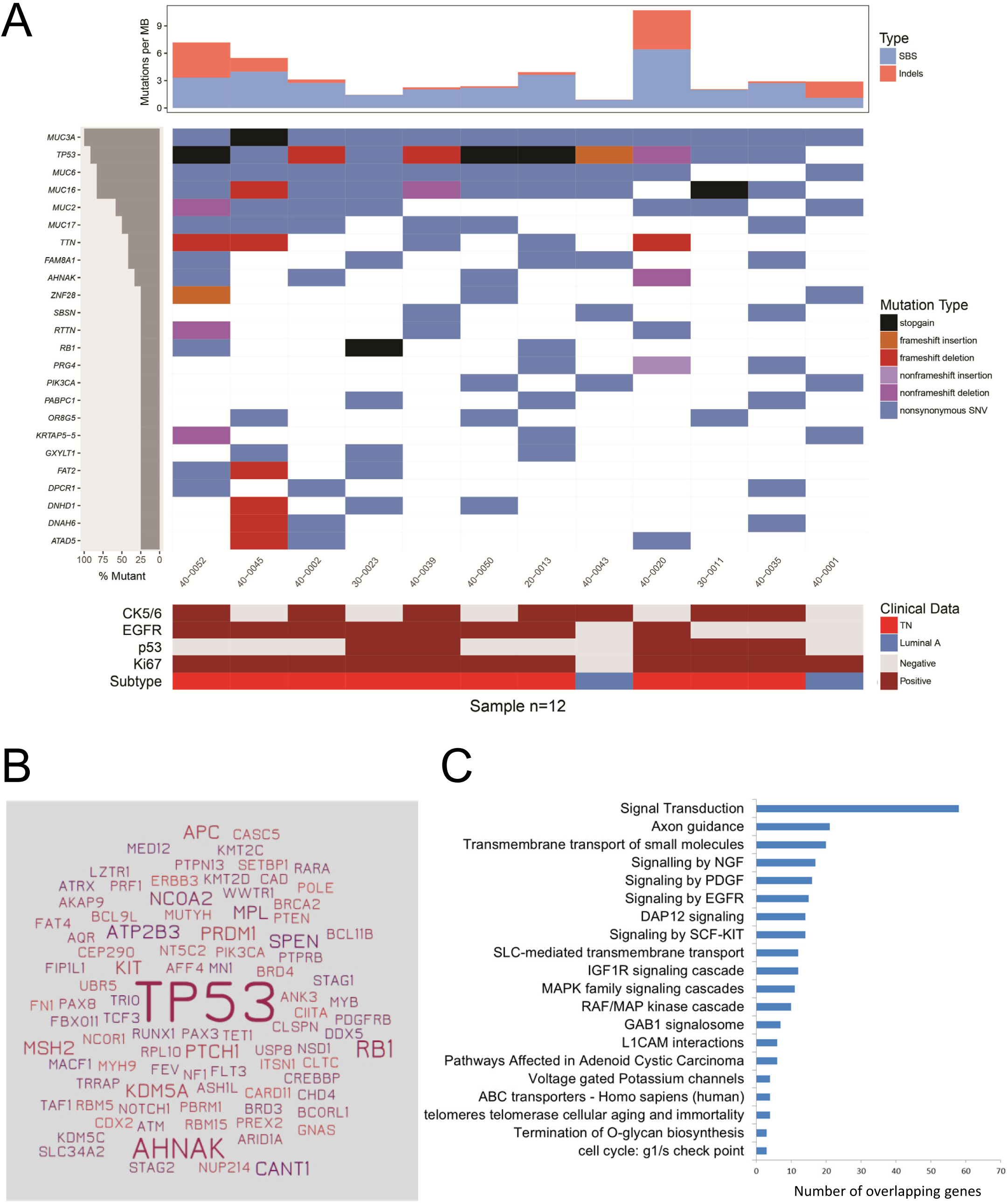
Whole exome sequencing results in 12 PRECAMA samples. Only coding non-silent somatic mutations are considered. (A) Mutation rates (top panel), top mutated genes and their mutation types (middle panel), and IHC features (lower panel), sorted by top mutated genes. Luminal A: ER+/HER2-; Triple Negative: ER-/PR-/HER2-. (B) All cancer genes somatically mutated in the 10 TN samples are depicted, the size of gene names being proportional to the number of sample mutated for each gene. (C) Pathways enriched (q-value <0.05) in the list of genes mutated in TN cases with AF>20% and predicted deleterious/probably deleterious by Polyphen (N=333 genes). Number of overlapping genes in each pathway is shown.

### Mutation patterns and signatures in premenopausal BC

To study the underlying mutational processes involved in the development of preBC tumors in the studied populations, we analyzed somatic mutation patterns in the *TP53* gene and at the exome-wide level. As shown in **Fig 3A**, the distribution of *TP53* mutation substitution types in PRECAMA samples was different from the one observed in other datasets of women younger than 45 years. There was in particular a higher proportion of G:C>T:A mutations in PRECAMA samples compared to BC tumors from young women from the IARC TP53 Database (p-value = 0.004; Chi-square test) or TCGA/METABRIC (p-value = 0.05; Chi-square test) datasets. In fact, G:C>T:A was the most frequent type followed by indels, while G:C>A:T at CpG was the most frequent type in the other datasets. *TP53* G:C>T:A mutations were observed in all subtypes while indels were more frequent in TN cases (**Fig 3B**). *TP53* indels were truncating mutations (predicted to result in loss of p53 protein expression) in 6/10 cases, and 5/6 of these truncating mutations were indeed associated with null p53 IHC staining (see **S1 Table**). Thus, while the presence of frequent *TP53* truncating mutations in TN subtype was similar to previous reports [26], the overall high frequency of G:C>T:A mutations was unexpected.

**Fig 3.**
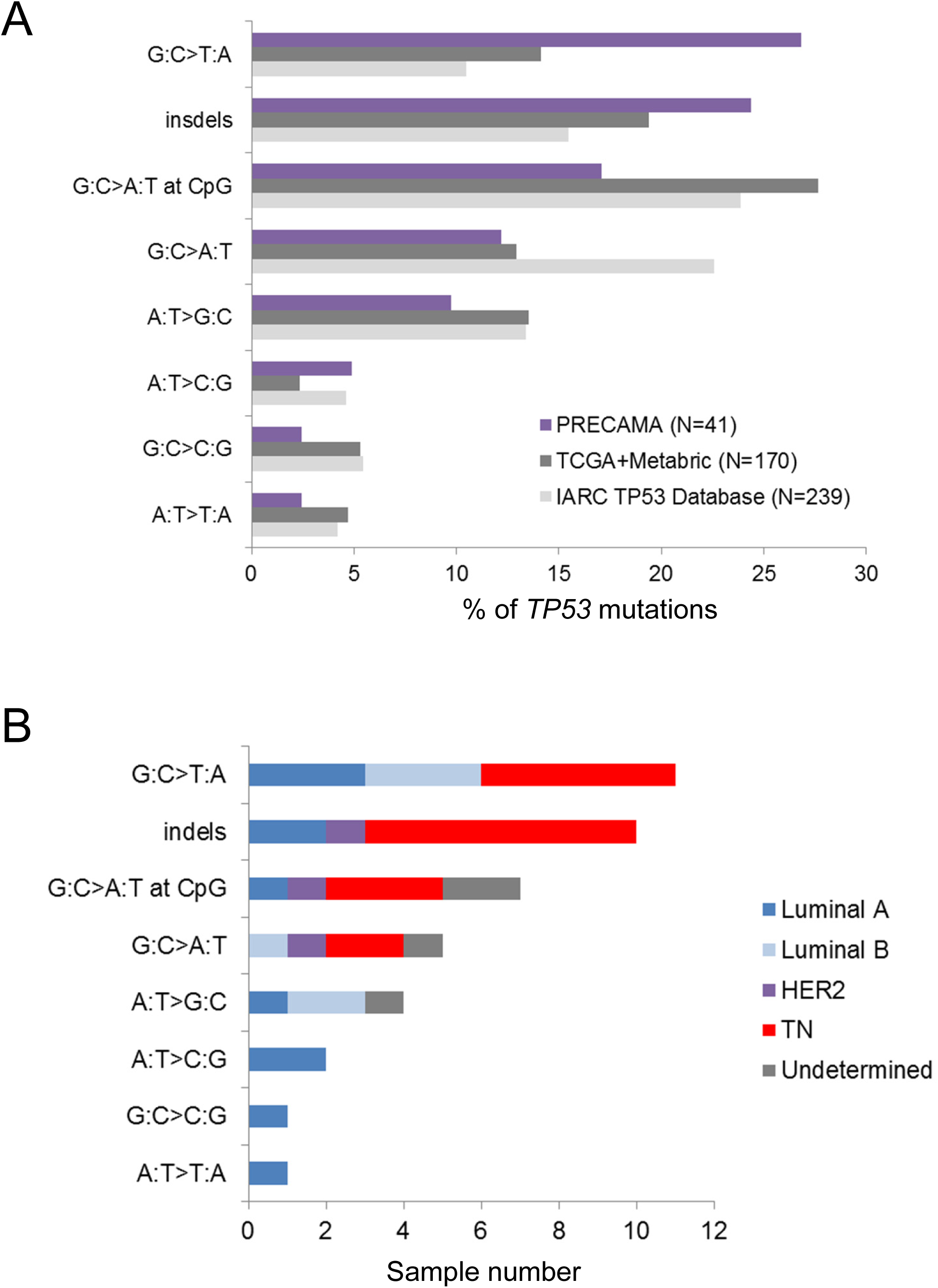
*TP53* mutation type distributions in preBC. (A) Distribution of mutation types in PRECAMA samples are compared to those observed in datasets of women <45 years old selected from the TCGA and METABRIC series [16, 17] or the IARC TP53 Database [18]. (B) IHC subtype distributions of PRECAMA samples in each mutation type category. Luminal A: ER+/HER2-; Luminal B: ER+/HER2+; HER2 Enriched: ER-/HER2+; Triple Negative: ER-/PR-/HER2-.

Mutational signatures at the exome-wide level were analysed using a dataset including the 12 PRECAMA samples and 453 BC samples from TCGA (see Materials and Methods). We identified 6 signatures that matched with previously reported signatures (**Fig 4A and S6 Table**). The estimated contribution of each signature to the mutation load in PRECAMA samples (**Fig 4B**) showed that 5/6 signatures had a contribution above 20% in at least one sample. Sig.A had the highest median contribution in these samples (24.3%). Sig.A matched with COSMIC signature-3 that has been established as a biomarker of homologous recombination defects through genetic and epigenetic inactivation of *BRCA1/2* pathway, a distinctive feature of basal-like tumors [7, 27, 28]. Sig.B, contributing in 6/12 samples, matched with signature 26, proposed to be linked to defective DNA repair and previously reported in breast cancer. Sig.C, contributing in 6/12 samples, matched with several signatures characterized by C>T mutations outside CpG sites, including experimental signatures induced by alkylating agents (MNNG and MNU) in rodent systems [29, 30], COSMIC signature-11 observed in recurrent brain tumors of patients treated with MNNG [31], and COSMIC signature-30 of unknown origin but previously observed in some BC. Sig.D, that matched with COSMIC signature-18, was mainly observed in one sample where it contributed to 90% of the mutation load and where the overall mutation load was the highest. The origin of this signature in BC remains to be established but it has recently been associated with germline mutation in the repair enzyme *MUTYH* in colorectal and adrenocortical carcinomas [32, 33]. Interestingly, the sample in which Sig.D dominated carried a truncating somatic mutation in *MUTYH* (see **S1 Table**). Finally, Sig.E, characterized by C>T mutations at CpG site and matching with COSMIC signature-1 known to be due to spontaneous deamination of 5-methylcytosine (also referred as the “age” signature), had significant contribution in only 3 samples.

**Fig 4.**
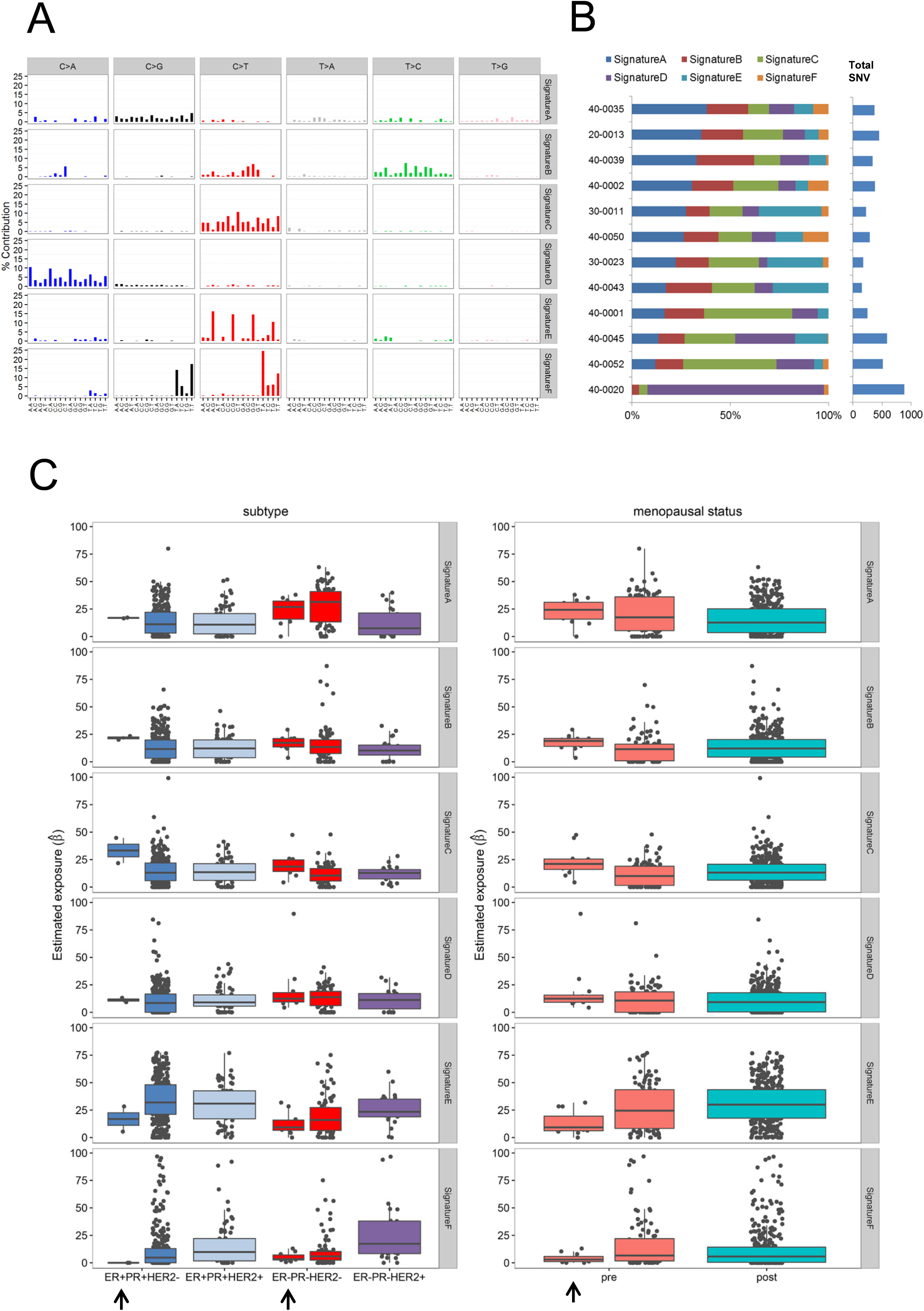
Mutational signatures identified in TCGA and PRECAMA samples, and their relationship with tumor subtype and patient menopausal status. (A) The six mutational signatures identified in 453 TCGA plus 12 PRECAMA samples. The six types of base substitutions are color-coded and further stratified by their adjacent 5′ and 3′ sequence context. Sig.A matches with COSMIC signature-3; Sig.B matches with COSMIC signature-26; Sig.C matches with COSMIC signatures-11/19/23/30 and experimental signatures of MNU and MNNG; Sig.D matches with COSMIC signature-18; Sig.E matches with COSMIC signature-1; Sig.F matches with COSMIC signatures-2/13 (see Supplementary Table 1, Additional File 1). (B) Percent contributions of the six mutational signatures to the SNVs found in PRECAMA samples. (C) Percent contributions of the six mutational signatures in PRECAMA and TCGA samples stratified by tumor IHC subtypes (left graphs) and by menopausal status (right graphs). PRECAMA samples are indicated by an arrow.

In **Fig 4C**, we explored possible systematic differences of signature contributions between IHC subtypes, and by menopausal status or study source (TCGA vs PRECAMA). Sig.F that matched with COSMIC signature-2 and COSMIC signature-13, linked to mutagenesis by APOBEC, was more prevalent in HER2-enriched subtype cases and underrepresented in TN cases (p-value < 5E-4), as reported before [34]. This fits with the fact that we do not find a significant contribution of APOBEC signature in the PRECAMA samples (median contribution of 2.9%) as we only analysed TN and Luminal A cases. COSMIC signature-3 (Sig.A) was enriched in TN cases (p-value < 5E-4) and preBC (p-value = 0.03). This signature was the predominant one in PRECAMA TN cases (median contribution of 26.8%). The contribution of the “age” signature (Sig.E) was lower in TN cases than in all other subtypes (p-value < 5E-4) and also lower in preBC compared to postBC (p-value = 0.006). It was the lowest in PRECAMA samples. The contribution of Sig.D was slightly higher in TN cases compared to all other subtypes (p-value = 0.01) and was the main contributor to the mutation load in one PRECAMA sample (**Fig 4B**). As this sample carried a somatic mutation in *MUTYH,* and a recent study found germline mutations in *MUTYH* in young women with breast cancer [35], it will be interesting to further study the role of *MUTYH* alteration in TN and premenopausal BC. In stratified analyses by IHC subtypes, signatures’ contributions in PRECAMA TN cases were similar to the ones observed in TCGA TN samples except for Sig.C (contribution was higher in PRECAMA than in TCGA TN samples, p-value = 0.006, median contributions: 18.6% vs 10.4%). Because Sig.C matched with several signatures, including signatures linked to exposure to alkylating agents not expected in these treatment naïve samples, its origin remains to be established. There was no effect of menopausal status on signature’s contributions when taking into account IHC subtype using linear models (all p-values > 0.16; permutation tests of linear regression model).

## Discussion

The results obtained in this pilot phase of the PRECAMA study demonstrate feasability regarding advanced genomic analyses of the tumor and blood samples collected at multiple sites in LA. They provide a preview of the molecular features of preBC in that population, with interesting mutational patterns that deserve further study.

Indeed, over 92% of samples processed for IHC analyses were successfully scored for 7 markers (only 20/253 were excluded due to absence of invasive tumor or to insufficient tissue for testing), and 80% of samples processed for DNA extraction yielded DNA quantities and quality compatible with genomic analyses (136/172 samples yielded more than 200 ng of DNA). With a target of 1500 cases recruited for the full study (with Guatemala and Brazil joining the study), this will be the largest series of preBC in LA women where genomic characterization of the tumor will be performed.

The IHC analyses showed a majority of ER positive cases and a proportion of TN subtype similar to previous reports in Hispanic women [36]. The overall prevalence of ER negative tumors in PRECAMA was substantiated by sequencing results on the 8-genes panel analyzed here. Indeed, *TP53* mutations, that are strongly associated with ER negative status [37, 38], were found in 32% of the cases, fitting with an overall 28% of ER negative cases. Also, the frequency of *AKT1* mutations, typical of ER positive cases [17, 39], was higher in PRECAMA than in the comparative dataset of young women. Continued enrollment will allow us to determine more precise estimates of subtype distribution in the LA population and to explore potential differences in tumor subtype distributions between countries.

Although overall tumor characteristics were more similar to those described in postBC than preBC, IHC staining with Ki67 showed high levels of staining in these preBC, even in Luminal A cases (72% positive cases), which is consistent with previous reports on preBC [40, 41]. Liao et al,. 2015 [42], recently compared the molecular features of preBC versus postBC from TCGA and METABRIC datasets using multi-omic data integration. They reported no difference in gene expression between preBC and postBC in ER negative cases but significant differences in ER positive cases, with activation of integrin signaling and EGFR pathways and TGFβ as the top upstream regulator in preBC. It would thus be important in future studies to assess whether activation of these pathways drives the level of proliferation reflected by high Ki67 positivity in ER positive preBC as they may be potential clinical targets.

The characteristics of the mutations found by target sequencing of the 8-gene panel was similar to the ones observed in other series of BC, with classical hotpots found in *AKT1* and *PIK3CA*, a majority of missense mutations found in *TP53*, a higher proportion of truncating *TP53* mutations in TN cases compared to other subtypes, and an expected distribution of mutated genes within IHC subtypes. However, an interesting difference in the distribution of *TP53* single base substitutions was observed. The most frequent *TP53* mutation type was G:C>T:A that represented 27% of all *TP53* mutations, a figure close to twice the one observed in the comparative datasets used here, and that matches those reported in lung cancers linked to PAH exposure [43, 44]. This pattern is thus unexpected in breast cancer. These G:C>T:A mutations do not exhibit a strand bias, do not cluster at any hotspot, and seemed similarly distributed within IHC subtypes or country of origin, although numbers are still too low to draw any conclusion. As these results may suggest a specific, yet unknown, mutational process at the origin of *TP53* mutations, it will be important to confirm them in the full study.

Exome-wide mutation profiling of a subset of basal-like TN tumors confirmed that *TP53* and *RB1* were the only cancer genes recurrently affected by deleterious mutations (>2 samples). These results are concordant with previous reports on TNBC of basal-like type that showed predominance of *TP53* mutations and of *TP53* and *RB1* pathways alterations [39, 45]. These reports also suggest activation of the PIK3CA/AKT pathway based on gene copy number analyses (*PIK3CA* gene amplification, *PTEN* gene deletion) and protein phosphorylation assays [39]. Here we found one activating *PIK3CA* mutation in 10 TNBC samples, which is in the range of previous reports (9%). However, as we limited our analyses to SNVs and small indels, we could not further assess the functionality of the *PIK3CA* pathway. Pathway analysis of potentially functional mutations across all genes showed enrichment of signal transduction pathways including EGFR, PDGF and IGF1R, and mutational signatures showed a large contribution of DNA repair defects to the mutation load. These overall results on TN cases fit with our previous analyses of another series of TN cases from Mexico where transcriptomics analyses showed an overexpression of growth-promoting signals (including *EGFR*, *PDGFR* and *PIK3CA*), a repression of cell cycle control pathways (*TP53*, *RB1*) and a deregulation of DNA-repair pathways [46].

Our exploratory analysis of exome-wide mutational signatures in relation to IHC subtype and menopausal status in TCGA and PRECAMA samples showed that the contributions of mutational signatures are determined by the tumor subtype but not the menopausal status, and that PRECAMA TN cases showed contributions similar to TCGA TN samples for 5/6 signatures identified in the analyzed set.

Some limitations of the results presented should be noted. First, the prevalence of IHC subtypes are based on still limited numbers and may thus not be representative of the distribution at the population level. Second, HER2 status confirmation by FISH could not be done in this pilot phase and thus the prevalence of Luminal B or HER2-enriched subtypes may be under- or over-estimated. Third, the exome data have been performed on a limited number of cases to establish feasability of these assays. Results on this small set did show feasability and allowed us to identify both similarity and differences in genomic alterations compared to other series of BC. Analysis of the full series will determine if any specific genomic feature may characterize preBC in LA women.

## Conclusions

This pilot results on PRECAMA tumors gives a preview of the molecular features of premenopausal BC in LA. Although, the overall mutation burden was as expected from data in other populations, mutational patterns observed in *TP53* suggested possible differences in mutagenic processes giving rise to these tumors compared to other populations. Further omics analyses of a larger number of PRECAMA cases in the near future will allow investigating relationships between these molecular features and etiological factors.

## Acknowledgements

The PRECAMA team includes International Agency for Research on Cancer (Coordinating Center) Investigators and Staff: Carine Biessy, Magali Olivier, Sabina Rinaldi, Isabelle Romieu; Investigators and Staff in Chile: Eva Bustamante, Fancy Gaete, Maria Luisa Garmendia, Jose Soto; Investigators and Staff in Colombia: Alberto Angel, Carlos Andres Ossa, William H. Arias, Gabriel Bedoya, Mauricio Borrero, Alicia Cock-Rada, Israel Díaz-Yunez, Carolina Echeverri, Fernando Herazo, Angel Hernández, Roberto Jaramillo, Edgar Navarro, Yorlany Rodas Cortes, Gloria I. Sanchez; Investigators and Staff in Costa Rica: Bernal Cortes, Paula Gonzalez, Diego Guillen, Viviana Loría, Rebecca Ocampo, Carolina Porras, Ana Cecilia Rodriguez; Investigators and Staff in Mexico: Gabriela Torres-Meija, Angelica Angeles-Llerenas, Jenny Tejeda; Investigators and Staff in Seattle: Peggy Porter.

The author wish to thank the substantial support provided by the research nurses and health workers as well as Tracy Lignini, Mathilde His, Dacia Cristin, Cecile Le Duc, Jordi de Battle, Talita Duarte-Salles, Ana Cristiana Ocampo.

The authors also wish to thank the women participating in the project for their time and commitment.

## Supporting information

**S1 File. S1-S6 Tables.** S1 Table: Demographics and molecular characteristics of cases analysed by NGS; S2 Table: Whole exome sequencing data metrics; S3 Table: Mutations in coding regions from whole exome sequencing and mutation calling with Strelka; S4 Table: List of mutated genes in TN cases with mutations present at an allele frequency above 20% and predicted to impact protein function (splice, truncating and non-synonymous predicted deleterious/probably deleterious by Polyphen-2) (N=333); S5 Table: Pathway analysis of 333 altered genes in TN samples; S6 Table: Cosine similarity values for the comparisons between each of the 6 extracted signatures and 37 published signatures.

**S1 Fig. IHC subtypes distribution in PRECAMA samples and in preBC extracted from METABRIC and TCGA studies.** Comparison of the distribution of IHC subtypes observed in PRECAMA and in preBC from a dataset extracted from METABRIC and TCGA (see Material and Methods). Luminal A: ER+/HER2-; Luminal B: ER+/HER2+; HER2 Enriched: ER-/HER2+; Triple Negative: ER-/PR-/HER2-.

**S1 Fig.**
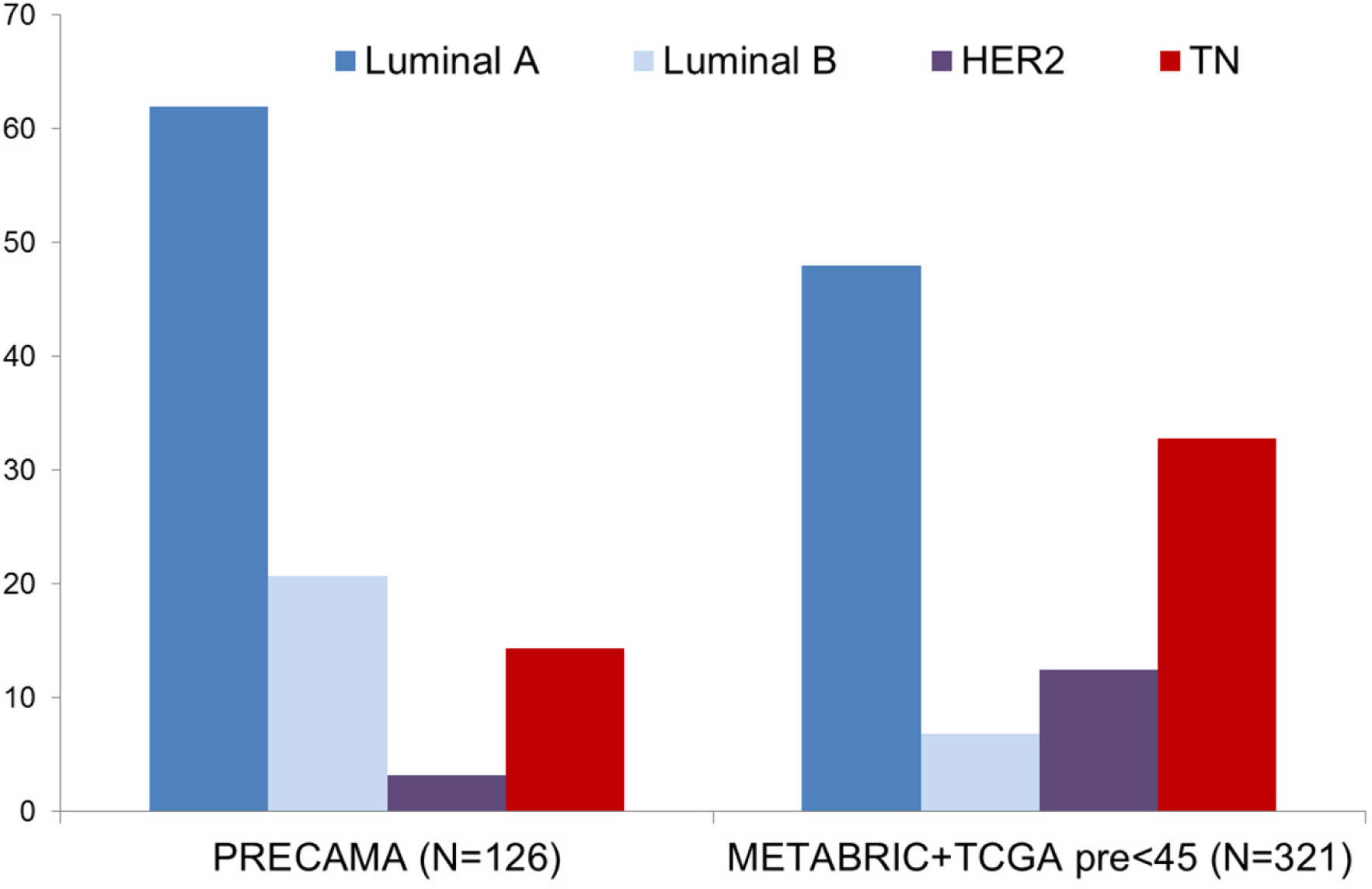
Comparison of the distribution of IHC subtypes observed in PRECAMA and in preBC from a dataset extracted from METABRIC and TCGA (see Material and Methods). Luminal A: ER+/HER2-; Luminal B: ER+/HER2+; HER2 Enriched: ER-/HER2+; Triple Negative: ER-/PR-/HER2-.

## List of abbreviations

BC: breast cancer
preBC: premenauposal breast cancer
postBC: postmenauposal breast cancer
TN: triple negative
TNBC: triple negative breast cancer
ER: estogen receptor
PR: progesterone receptor
WES: whole exome sequencing
IHC: immunohistochemistry
HR: homologous recombination
DMR: DNA mismatch repair

